# Boosting the analysis of protein interfaces with Multiple Interface String Alignment: *illustration on the spikes of coronaviruses*

**DOI:** 10.1101/2020.09.03.281600

**Authors:** S. Bereux, B. Delmas, F. Cazals

## Abstract

We introduce *Multiple Interface String Alignment* (MISA), a visualization tool to display coherently various sequence and structure based statistics at protein-protein interfaces (SSE elements, buried surface area, ΔASA, B factor values, etc). The amino-acids supporting these annotations are obtained from Voronoi interface models. The benefit of MISA is to collate annotated sequences of (homologous) chains found in different biological contexts *i.e.* bound with different partners or unbound. The aggregated views MISA/SSE, MISA/BSA, MISA/Δ ASAetc make it trivial to identify commonalities and differences between chains, to infer key interface residues, and to understand where conformational changes occur upon binding. As such, they should prove of key relevance for knowledge based annotations of protein databases such as the Protein Data Bank.

Illustrations are provided on the receptor binding domain (RBD) of coronaviruses, in complex with their cognate partner or (neutralizing) antibodies. MISA computed with a minimal number of structures complement and enrich findings previously reported.

The corresponding package is available from the Structural Bioinformatics Library (http://sbl.inria.fr)

## 1 Introduction

### Protein complexes and their interface

Understanding the stability and the specificity of protein interactions is a fundamental problem [1], both to unveil complex biological mechanisms and design therapeutics. From the thermodynamic standpoint, the binding affinity or equivalently the dissociation free energy, boils down to compute the ratio of three so-called partition functions [2]. These partition functions are themselves defined from the potential energy of the system, which in particular encodes interaction energies at play between atoms (electrostatic interactions, non covalent interactions, bonded interactions). This observation does not settle the question though, as the huge dimensionality of the system involving the biomolecules and the solvent precludes calculations, except maybe for small systems using massive calculations [3, 4].

This observation also explains the success of geometric models based on distances, surfaces and volumes, which originate with the seminal work of Richards for individual molecules [5] and Janin for complexes [6]. Such models proved instrumental to understand the specificity and affinity of interactions [1]; in some cases, they even proved good enough to estimate binding energies up to (of the order of) 1.4kcal/mol [7]. To study two partners forming a complex, such models require identifying the atoms in contact. To this end, Voronoi models, be their affine [8] or curved [9], provide a parameter free approach to model interfaces, and affine models in particular give a direct access to various structural parameters including the number of interface atoms and residues, the buried surface area, the number of binding patches, or the interface curvature [8].

These parameters are typically sufficient to compare interfaces for cases belonging to various protein families (protease-inhibitors, enzyme-substrate, antibody-antigen, signal transduction, etc) [10], or cases within a protein family [11]. On the other hand, in processing complexes specifically addressing a biological case, say a family of homologous proteins targeting the same cognate partner, it is also critical to study correlations between structural parameters describing interfaces and sequence related pieces of information. For sequences, the natural route in to resort to multiple sequence alignments (MSA) [12].

### Multiple Interface String Alignment (MISA)

In this work, we introduce MISAs, a visualization tool to display coherently various sequence and structure based statistics at protein-protein interfaces (SSE elements, buried surface area, ΔASA, B factor values, etc). The benefit of MISAs is to collate annotated sequences of (homologous) chains found in different biological contexts (unbound or bound with different partners).

We illustrate the interest to analyse the receptor-binding domain (RBD) or the virus SARS-Cov-2 causing the covid19 outbreak. First, we compare the receptor-binding domains (RBD) of SARS-Cov-1 and SARS-Cov-2 (two phylogenetically-related betacoronavirus). Second, we compare the interfaces of their RBD bound to their common cell receptor and with antibodies.

The implementation of MISAs available in the SBL (http://sbl.inria.fr, [13]).

## 2 Methods: MISAs – multiple interface string alignments

### 2.1 MISA

We assume a collection of crystal structures of the bound and unbound types. We call a polypeptide chain in these structures a *chain instance*. We also assume that a multiple sequence alignment is available for the sequences of the instances of interest. Practically, we expect two settings: (i) chain instances with identical sequence: since numberings used in the respective PDB files are identical, the identity alignment is used; (ii) homologous proteins: a multiple sequence alignment providing a coherent numbering of amino acids (a.a.) is taken for granted.

The terminology used in the sequel is as follows. We study chain instances and compare them against one another using one *MISA* for each so-called *MISA id*. A MISA is based on *interface strings* (i-string), which following conventions for sequence alignments, use the one-letter code of (a.a.) plus three additional symbols {_ ∗ −}. We now detail these concepts.

#### MISA id

A complex *C_i_* is specified by two sorted lists of chains ids *i.e. C_i_* = ({*A_j_*}, {*B_j_*}) for the two partners; each list is endowed with a partner/structure name. Likewise, an unbound structure is specified by a sorted list of chain ids *U_i_* = {*A_j_*}, and is also given a partner/structure name (*e.g.* antibody, antigen, etc.) We consider a collection of complexes 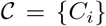, and optionally a collection of unbound structures 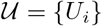. The minimal setup is naturally that of a single complex without any unbound structure.

Our goal is two build one MISA for each so-called MISA id, which we formally define as:

##### Definition. 1

*The* MISA id *of a chain in a complex or unbound structure is the string defined by the structure name followed by the index of the chain in its sorted list {A_j_} or {B_j_}. (Nb: by convention, indices start at 0.)*

Note that because of the sortedness, a given chain gets the same MISA id in a complex or unbound structure.

##### Example 1

*Consider two antibody-antigen complexes. C*_1_ = ({*H, L}, {A}*), *C*_2_ = ({*M, N*}, {*B*}), *each involving the heavy chains (chains H and M), the light chains (chains L and N), and the antigen (chains A and B). The two structure names are thus antibody (IG for short) and antigen (Ag for short). The MISA id of the heavy chain is IG_0, that of the light chain IG_1, and that of the antigen Ag_0.*

#### Voronoi interface in a binary complex

The two lists of chains in the specification of a complex *C_i_* define the so-called partners in the complex, called *A* and *B* for short. In the sequel, we build the Voronoi interface for these partners, using the Voronoi (power) diagram of the solvent accessible model [8, 1, 14]. More precisely, recall that the Voronoi model identifies pairs of atoms, one on each partner, which are either directly in contact as their Voronoi cells are neighbor, or are contacting a common crystallographic water molecule. Therefore, an a.a. contributing at least one interface atom is called an interface residue/a.a‥

Practically, we use the `sbl-intervor-ABW-atomic.exe` executable from the package Space_filling_model of the Structural Bioinformatics Library ([13], http://sbl.inria.fr).

#### Interface strings (i-strings)

In the sequel, we present MISAs informally, referring the reader to SI Section *Supplemental: formal specification of the method*. We represent each chain instance with a so-called *interface string* encoding properties of its interface amino acids. The interface string of a chain instance is a character string with one character per residue, and is actually defined from all instances with the same MISA id. To build this character string, we first build a *consensus interface* based on all Voronoi interfaces of all complexes in the set 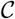. At each position of the consensus interface, the most frequent residue observed in all the bound structures is chosen as the consensus residue–with ties broken using the alphabetical order of the one letter code of a.a‥ Using this consensus interface, the residues of a given chain instance (bound or unbound) are assembled into the so-called interface string:

##### Definition. 2

*The* interface string *(i-string) of a chain instance is the string with one character per amino acid, defined as follows:*

1. *Residue not part of the consensus interface:*

- *Displayed with a dash “-” if it is part of the crystal structure, and underscore “_” otherwise.*
2. *Residue part of the consensus interface:*

- *Residue not found in the crystal structure: displayed with the star ‘*’.*
- *Residue found at the interface for this particular chain: displayed with the uppercase one letter code if the a.a. matches the consensus a.a., or with the lowercase letter otherwise.*
- *Residue not found at the interface for this particular chain, even though the corresponding position contributes to the consensus interface (for other chains): displayed in an italicized uppercase/lowercase letter. (Note that this is the case for all residues of unbound structures–as no partner implies no interface.)*

Practically, it is often sound to restrict the study to a range of amino-acids on the chains studied, whence the following:

##### Definition. 3

*A* window series *for a MISA id is a sequence of integer ranges of residue ids* {(*B_i_, E_i_*)} *used to restrict the display of the i-string.*

#### MISA

We use the MSA to align the symbols of interface strings, yielding:

##### Definition. 4

*The* MISA *of all chain instances in 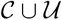 with the same MISA id is the multiple alignment of their interface strings*.

### 2.2 Colored MISAs

We further color the letters of a MISA to encode properties of biological/biophysical interest.

#### MISA/SSE: coloring based on Secondary Structure Elements

The type of SSE a given a.a. belongs to is especially useful when comparing bound and unbound structures, to assess perturbations in the hydrogen bonding network. Practically, we use the dictionary of SSE from [15].

#### MISA/BSA: coloring based on Buried Surface Area

Consider a chain instance in a complex. The BSA of this chain is defined as the accessible surface area (ASA) [5] of this chain in the partner alone minus the ASA of the chain in the complex. We report the BSA on a per residue basis, computed using the algorithm from [16].

##### Remark 1

*The Voronoi interface model identifies* privileged *contacts between atoms of the partners. Interestingly, selected interface atoms, generally backbone atoms, can be buried within their own subunit [17]. Since such atoms do not have any BSA (they do not have ASA in their own subunit), a residue containing only such atoms is an interface residue for which the BSA information is irrelevant.*

#### MISA/Δ ASA: coloring based on the variation of accessible surface area

A limitation of the BSA is that its calculation uses the geometry of the bound structure only. The calculation is thus oblivious to conformational changes which may be at play in case of induced fit or conformer selection. To mitigate the previous plot, we also provide the so-called Δ ASAcoloring scheme.

Consider an interface partner *P* (= *A* or *B*), in the complex, and consider the i-th residue of one of its chains. Let 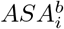 be the ASA of this i-th residue in the structure involving only the chains defining partner *P*. Also denote 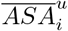 the average ASA of the i-th residue in unbound structures with the same MISA id. We compute for the i-th residue of a bound structure the quantity 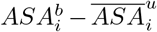 and display it with a color map.

##### Remark 2

*The quantity 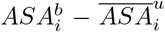 is undefined when the corresponding residues are absent in the unbound crystal structures (* in i-string).*

#### MISA/B-factor: coloring based on B-factors

The B-factors reflects the atomic thermal motions. Selected recent crystal structures report this information as a 3×3 ANISOU matrix (the anisotropic B-factor). In order to have a single quantity for all the crystals, the ANISOU matrix *B_anisou_* is converted into B-factor thanks to the formula *B_factor_* = trace(*B_anisou_*)/3 [18]. Optionally, one can choose to normalize the B-factor with respect to (i) all the residues in the chain, or (ii) all the displayed residues.

### 2.3 Availability and practical matters

MISAs are implemented within the Multiple_interface_string_alignment package of the Structural Bioinformatics Library ([13], http://sbl.inria.fr), see https://sbl.inria.fr/doc/Multiple_interface_string_alignment-user-manual.html.

The calculation and the analysis of MISAs is based on four scripts–see online documentation:

- `sbl-misa.py`: building MISAs from a description of complexes and possibly unbound structures.
- `sbl-misa-mix.py`: mixing selected colored MISAs into a single (html) file.
- `sbl-misa-bsa.py`: displaying the BSA of all (or selected user defined) residues.
- `sbl-misa-diff.py`: comparing i-strings and associated properties (in particular BSA) of two interfaces. Has two modes: comparing two i-strings; comparing one i-string and a list of interface residues typically coming from a publication.

Illustration of these scripts to investigate the questions discussed in the next section are provided in the jupyter notebook which accompanies our software.

## 3 Results: studying the receptor-binding domain (RBD) of coronaviruses

Similarly to other enveloped viruses, the SARS-Cov-2 virus enters cells using fusion proteins. Having recalled the known facts on this mechanism, we show that MISAs are helpful in two respects: (i) to automate such studies and refine some of the findings reported so far, and (ii) to provide a coherent view of several sequence-structure based statistics, which is instrumental in particular to understand the binding affinity of SARS-Cov-2 RBD for its target.

### 3.1 Biological problem

Coronavirus entry into cells is mediated by the so-called spike that is responsible for receptor binding and membrane fusion [19, 20]. The covid19 outbreak triggered an impressive number of structural studies on the entry of SARS-Cov-2, namely [21] [22] [23] [24] [25] [26] [27]. Interactions between the spikes of SARS-Cov-2 with antibodies have also studied [28],[29], [30]. These papers naturally unveiled critical aspects of this complex process, based on experiments and also sequence based and structure based analysis.

Spikes are homotrimers of the protein S [31, 32]. Each chain consists of two domains S1 and S2, separated by cleavage sites denoted S1/2 and S2’ [33, 27, 29]. Domain S1 contains the receptor-binding domain (RBD), while domain S2 contains the machinery responsible for the fusion of the virus envelope with the membrane of the cell. In short, the main steps of the mechanism are as follows [33, 23]: (1) Attachment of the RBD to its target thanks to the receptor binding-motif (RBM), (2) proteolysis cleavage/activation at S1/S2 removing the S1 subunit, (3) second cleavage at S2’, refolding of fusion machinery - anchoring of fusion peptide into the target membrane, (4) membrane fusion and genome delivery into the cytoplasm. In human, both SARS-Cov-1 and SARS-Cov-2 targets ACE2 [26], a membrane bound enzyme catalysing the hydrolysis of angiotensin II into angiotensin, as a receptor to enter into cells. The following facts are noticeable [23]. The virulence of SARS-Cov-2 owes in particular to the presence of a polybasic cleavage site (insertion of three R) collated to the usual S1/S2 cleavage site. The RBD of SARS-Cov-2 stands up less often than that of SARS-Cov-1, which likely favors evasion from the immune system. (Nb: the two conformations are called up/down or closed/open.) This also disfavors binding to the target (binding in the down configuration yields steric clashes), but the lesser proportion of RBD standing up is *rescued* by a higher binding affinity for ACE2 [23]. Interestingly, a phylogenetically distant human alphacoronavirus, HCoV-NL63 also targets ACE2 as a receptor.

In the following, we illustrate the ability of interface string and MISAs to shed light on these complex facts. In doing so, our goal is a presentation of the merits of MISAs, rather than an arbitration of the merits of different publications which investigated the same systems. For this reason, we use a minimal number of crystal structures to get informative MISAs, namely two bound and two unbound structures, choosing those with the highest resolution. As we shall see, even such a simple study sheds new light on a number of published facts.

### 3.2 Comparing the RBD of SARS-Cov-1 and SARS-Cov-2 in complex with ACE2

#### Baseline

The spikes of SARS-Cov-1 and SARS-Cov-2 are highly homologous proteins binding ACE2. Pairwise alignment shows that the overall percent protein sequence identity is about 76% [34]. The S1 subunit is less conserved (64% identity) than S2 (90% identity). Within S1, the receptor-binding domain (RBD) is 74% identical. The initial analysis of the complex between the RBM of SARS-Cov-1 and its target ACE2 was first carried out in [35]. In this work, it is noted that the BSA at the interface is ~ 1700Å^2^, and that the interface involves 18 a.a. on the receptor side and 14 a.a. on the spike side. These analysis are extended to SARS-Cov-2 and other SARS viruses, in particular in [25, SI Fig 1], [24, Fig 2B], and [27, Fig 2C]. These recent publications take the 14 a.a. identified in [35] for granted.

**Figure 1:**
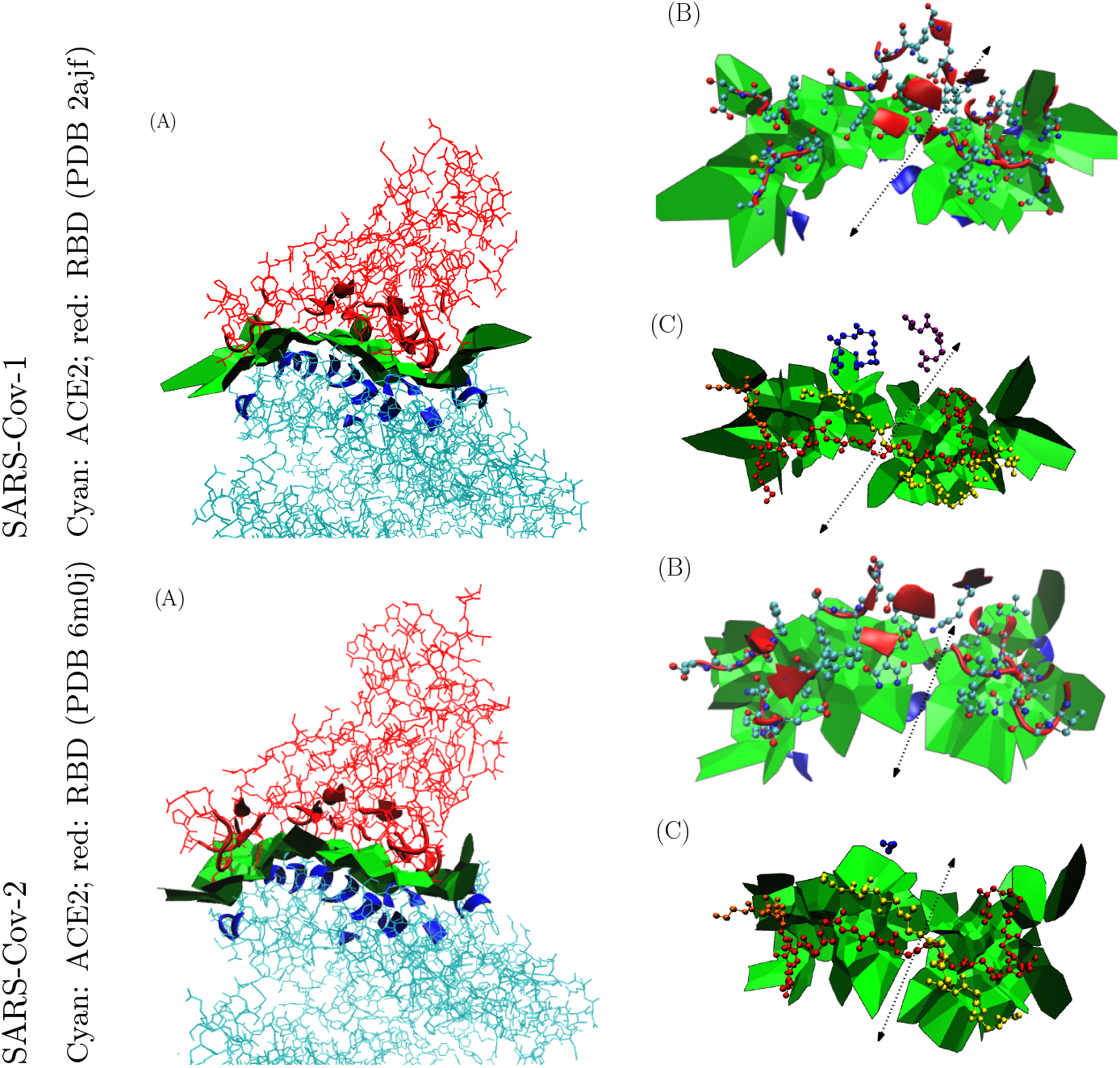
SARS-Cov-1 and SARS-Cov-2: Voronoi interfaces of the RBD in complex with ACE2. **(Panel A)** Side view of the Voronoi interface, with contributing a.a. as new ribbons. Note in particular the many disconnected contributions of SSE of the RBD to the interface. **(Panel B)** Top view of the interface, with interface a.a. as ribbons, and their side chains in CPK mode. **(Panel C)** The four or five ranges of consecutive a.a. identified by MISA, with backbone in CPK mode, on top of the Voronoi interface. SARS-Cov-1: 390-396 (violet), 404-408 (blue), 426-443 (yellow), 460-463 (orange), 470-492 (red); SARS-Cov-2: 417-421 (blue), 439-456 (yellow), 473-477 (orange), 484-506 (red)

**Figure 2:**
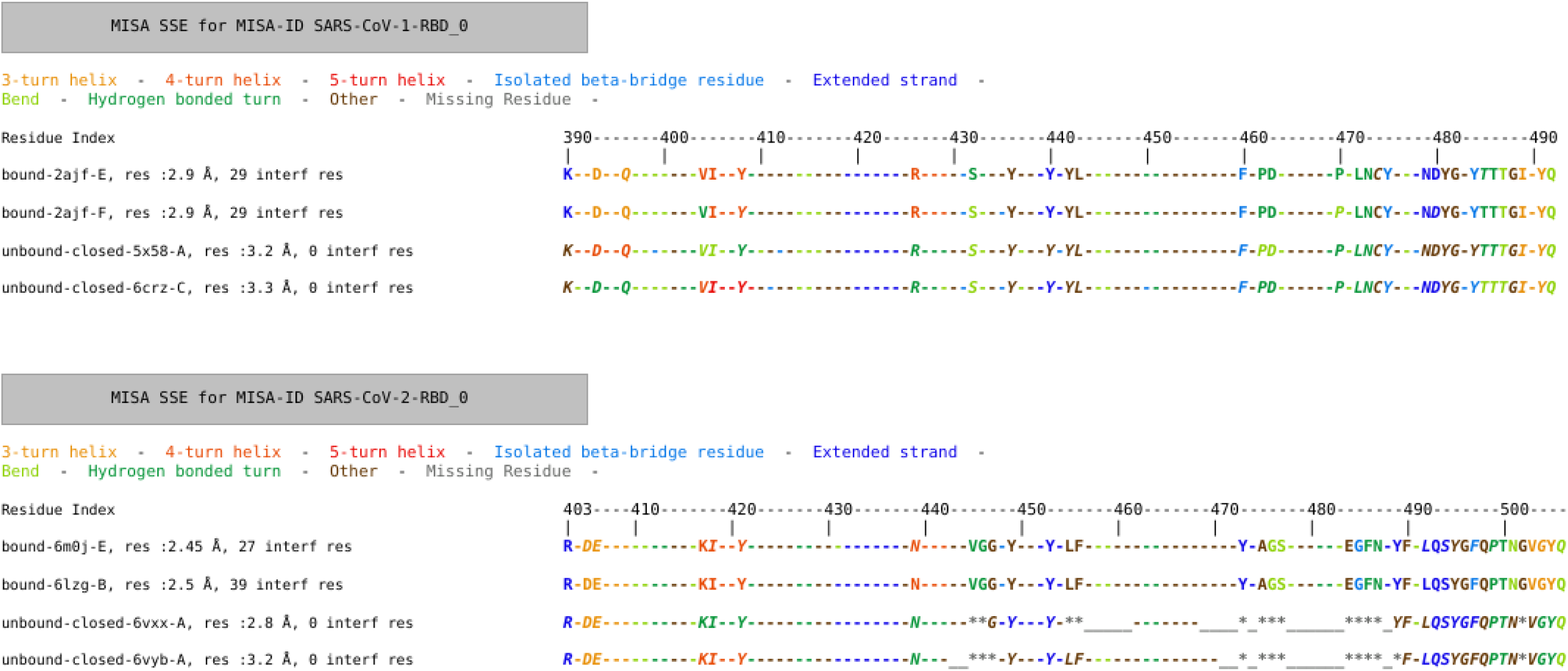
MISA/SSE for structures of Fig. 1.

Note that in furthering these analysis, we sometimes refer to the jupyter notebook which accompanies our software, and propose selected analysis using the four scripts of the Multiple_interface_string_alignment package.

#### Whole Voronoi interfaces

The Voronoi interface calculation yields comparable results (Figs. 1 and 2). For SARS-Cov-1 (PDB: 2ajf, RBD: chain E, ACE2: chain A), one obtains 246 interface atoms making up 2 binding patches, and defining overall 414 pairwise contacts. For SARS-Cov-2 (PDB 6m0j, RBD: chain E, ACE2: chain A), one gets 212 interface atoms also disposed on two binding patches, and defining overall 371 pairwise contacts. The two interface patches correspond to the transition between a beta sheet and loops on the RBD (Fig. 1(B,C)), dotted line-segments). Moving to interface strings (Fig. 2, we note that the RBD contribution is is characterized by five main stretches along the backbone. Interestingly, two stretches (yellow and red) span the whole interface, crossing on top of one another in a X like figure (Fig. 1(C)).

#### Buried surface area

The BSA is essentially proportional to the number of interface atoms [1], and is also a key parameter to estimate binding affinities [36, 7].

We first inspect BSA values obtained for two complexes in each case (SI Figs. 5 and 6), and compare them against the aforementioned 1700Å^2^: SARS-Cov-1: BSA(2ajf chains E+A)=925+888=1813Å^2^; BSA(2ajf, chains F+B)=864+817=1681 Å^2^; SARS-Cov-2: BSA(6m0j chains E+A)=887+843=1730Å^2^; BSA(6lzg, chains B+A)=1120+1098=2218Å^2^). While the values are coherent for SARS-Cov-1, it appears that the complex found in 6lzg features 30% more BSA. Inspecting the number of interface atoms readily provides the explanation (output of `sbl-intervor-ABW-atomic.exe`, see jupyter notebook): for 6lzg, one get 156 interface atoms for the RBD, 164 interface atoms for ACE2, and 22 water molecules sandwiched between the partners; these numbers drop to 101, 110 and 1 respectively for 6m0j. This reminds us a well known fact, namely that solvation by water molecules with low temperature factors (80 units in our case) may significantly change the interface model [8].

Beyond these global values, MISA/BSA gather BSA values for all atoms of the same a.a. (SI Fig. 5). Two interesting facts emerge. First, both for SARS-Cov-1 and SARS-Cov-2, the residues which bury the largest ASA are found at both endpoints of the fifth stretch (SI Fig. 5), contributing to the *stabilization* of the interface. For SARS-Cov-1, one notes in particular: Y475, T486, Y491; and for SARS-Cov-2: F486, Y489, T500, Y505. We also note that some of these a.a. bury a great deal of SAS (SI Fig 6), in particular Y475 for SARS-Cov-1, and Y489 for SARS-Cov-2.

#### Interface composition

We proceed with the inspection of the number of interface a.a., using the 14 a.a. reported in [35] as baseline.

Focusing on the RBD of SARS-Cov-1, we note that this number is precisely 29 for the crystal structures used (Fig. 2, MISA/SSE). It should be stressed that the this is not an artifact since two atoms are in contact in a Voronoi model provided that their Voronoi restrictions intersect. (Nb: the restriction of a ball is the intersection of the atomic ball and the Voronoi/power region.) We note in particular that three residues omitted from [35] have BSA larger than 30 Å^2^in (2ajf, chain E): L443: 34.93 Å^2^, P462: 49.79 Å^2^, and I489: 32.87 Å^2^(jupyter notebook).

Switching to SARS-Cov-2, these numbers are 27 and 39 respectively, for the aforementioned two crystal structures–a discrepancy owing to solvation by structural water. Letting these count alone, a merit of MISAs appears immediately: as opposed to previous presentations, which single out the 14 a.a. of interest in the whole RBM sequence, it appears immediately that the interface a.a. are located within five main stretches along the sequence.

Interface strings in MISAs also single out the relatively high abundance of tyrosine (Y), and glycine (G) amongst interacting residues. Such a.a. have been shown to cooperatively account for specific recognition in particular in antibody-antigen interactions [37]. The role of tyrosines at interfaces is well documented [38], both in terms on interaction types (non polar, cation-pi and H bonds) and dynamics (the rigid side chain entails a lesser entropic penalty upon binding). Again, this is not visible from previous representations [25, SI Fig 1], [24, Fig 2B], and [27, Fig 2C] since the interface amino acids are not identified in a systematic manner.

#### Conformational changes and binding affinity

MISA/SSE show that the RBD of SARS-Cov-2 is much less structured than that of SARS-Cov-1: while all residues are resolved in the unbound structures of SARS-Cov-1, a total of 12 of them are not so in the unbound structures of SARS-Cov-2 processed here (* in the i-strings, Fig. 2). The a.a. concerned are nonpolar (AGGF) or uncharged polar (NSY). Hydrophobic patches on protein surfaces are notorious and contribute to the recognition of cognate substrates [39]. The balance observed here hints at a rather hydrophobic patch also preserving the ability to make hydrogen bonds via NSY. The color changes in MISA/SSE convey the same information: more frequent color changes for SARS-Cov-2 show that binding triggers more significant changes of hydrogen bonding patterns.

The analysis of MISA/ΔASAcomplements this view (SI Fig. 6). For SARS-Cov-1, the interface a.a. have a balanced behavior, some gaining some loosing ASA, in the mild range (−10‥+30Å^2^). For SARS-Cov-2 on the other hand, the scale is shifted towards negative values (< −150‥ + 35Å^2^). In other words, the binding entails a compactification of the RBD in SARS-Cov-2.

Speaking of conformational changes, we further our analysis with a clustering of structures in terms of lRMSD, which is of special interest to assess the conformational changes between the up and down conformation of the RBD. In this case, no significant difference is observed in terms of lRMSD computed from all backbone atoms (SI Fig. 7). Phrased differently, MISAs show subtle details at the individual a.a. level, which are overlooked by global lRMSD calculation.

Concluding, the previous discussion shed lights on the determinants of the affinity of the RBDs of SARS-Cov-1 and SARS-Cov-2 for their receptors [23]. For SARS-Cov-2, the unstructured nature hints at a large entropic penalty upon binding. This penalty is clearly counterbalanced by a strong enthalpic component in the dissociation free energy, which we see in particular from the BSA.

### 3.3 Interactions between the RBD of SARS-Cov-2 and antibodies

Neutralizing antibodies targeting the spikes of viruses can implement various strategies [40], including (i) masking the RBD, (ii) preventing the conformational changes triggering the fusion envelope-membrane, or may be more surprisingly by (iii) mimicking the receptor and triggering early conformational changes of the fusion machinery far away from the cell membrane [28]. We illustrate again the interest of MISAs in this context.

We first consider the complex (SARS-Cov-2-RBD, IG CR3022), the latter being an IG isolated from a convalescent SARS patient. Since this IG also targets the RBD, two critical questions are investigated in [22]: first, the comparison of the complexes complexes SARS-Cov-1-RBD with CR3022 and SARS-Cov-2-RBD with CR3022, so as to understand in particular the lesser binding affinity of CR3022 for SARS-Cov-2-RBD (Kd=115nM versus Kd=1nM); second, the absence of in vitro neutralization of SARS-Cov-2 by CR3022, taking into in particular the up/down conformations of the RBD and the role of glycosylation.

Interestingly, the first point use MISAs prepared *by hand* [22, Fig. 2A]. The systematic nature our automatically generated MISA/SSE shows modest albeit interesting refinements. First, one sees immediately that the regions of the RBD targeted by ACE2 and CR3022 are disjoint (Figs. 3 and 4; jupyter notebook). Second, using the two complexes RBD-CR3022 in the asymmetric unit of the crystal, we note that the interface involves 33 rather than 28 a.a. which bury surface area. Those omitted from [22, Fig. 2A] are sitting on the boundary of the interface though (Fig. 3(B, VdW balls)). MISA/BSA and MISA/Δ ASAalso immediately identify those a.a. with the largest contribution to BSA (Figs. 8 and 9). Interestingly too, we note that binding with CR3022 triggers significant conformational changes with significant loss (Y380) or gain (H519) of ASA, as evidence by MISA/Δ ASA.

**Figure 3:**
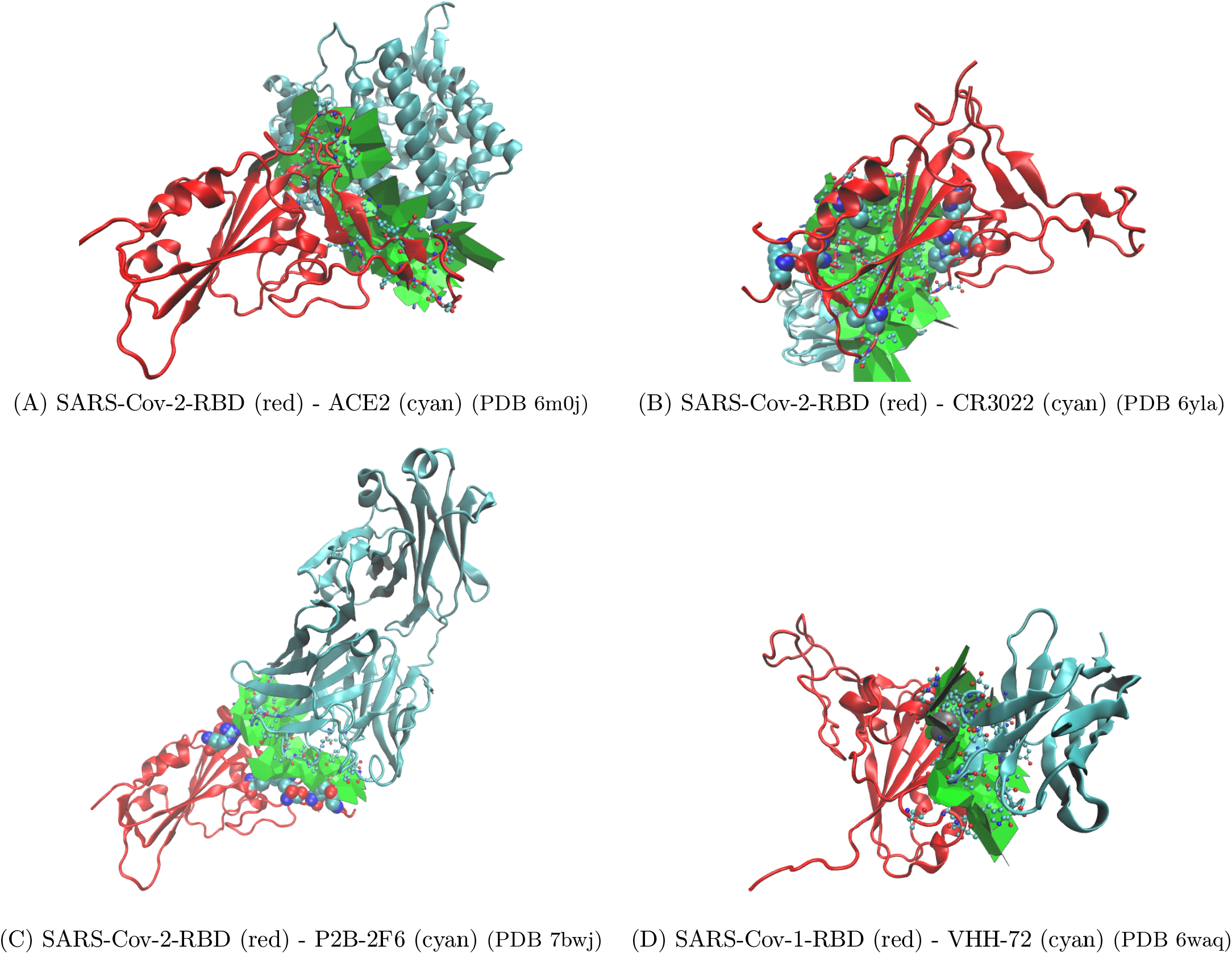
SARS-Cov-2 and SARS-Cov-1: RBD bound to the various IG / VHH: Voronoi interfaces. (A) Complex with ACE2 presented for reference (A). (B) Complex from [22]. Residues in VdW mode are identified by the Voronoi interface model but are not listed in [22]. (C) Complex from [30]. Residues in VdW mode are identified by the Voronoi interface model but are not listed in [30].

**Figure 4:**
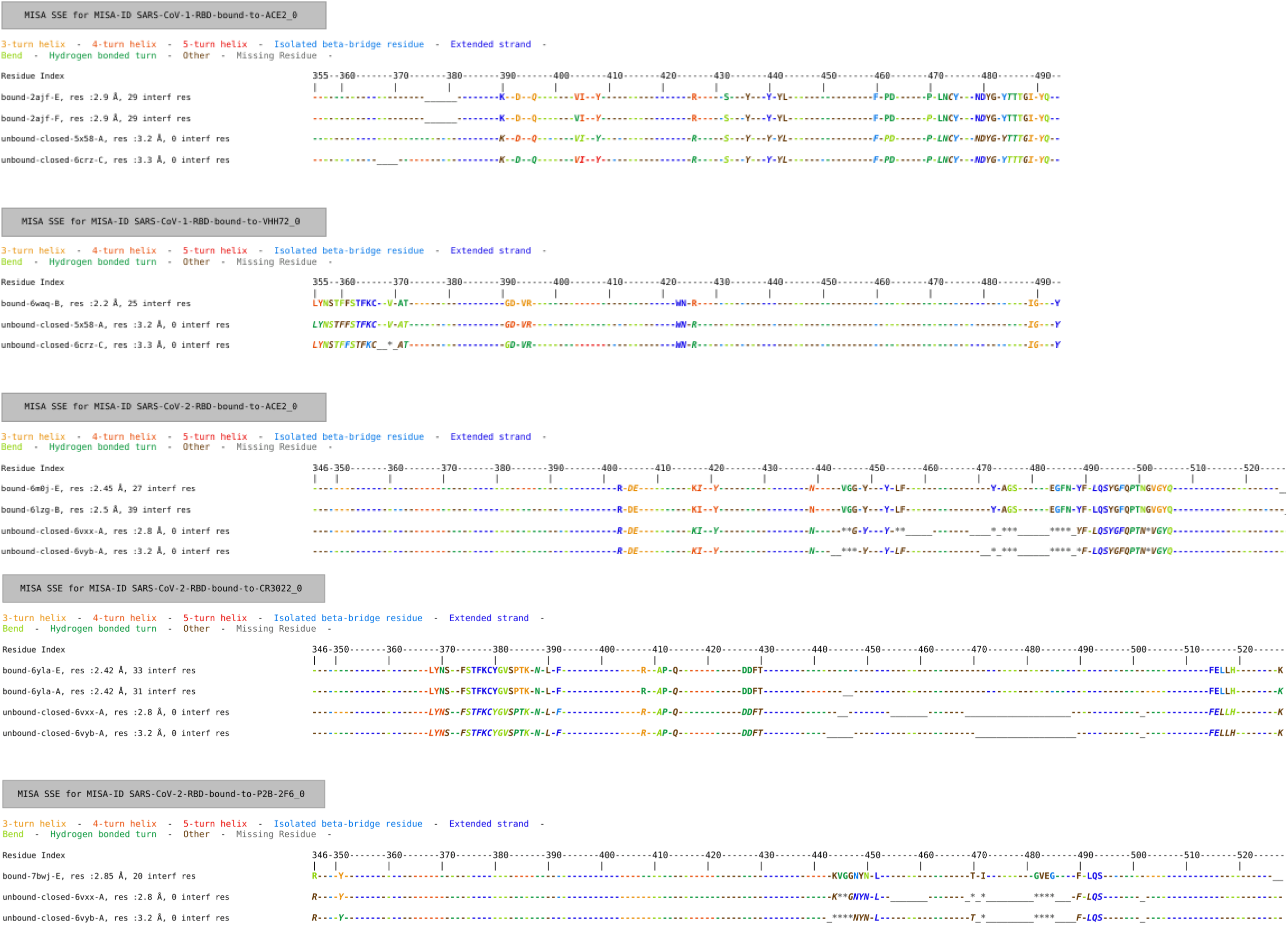
MISA/SSE for structures of Fig. 3. MISA of ACE2 not shown.

As a final illustration, we also compare the interface (SARS-Cov-2-RBD, ACE2) against that of (SARS-Cov-2-RBD, P2B-2F6) [30]. With 10 shared residues, 10 exclusive residues for RBD - P2B-2F6, and 29 exclusive residues for RBD - ACE2, this comparison shows a clear competition between the two binders (jupyter notebook). A direct comparison against the 12 residues reported in [30] also shows that 8 residues reported by the Voronoi model have been omitted (Fig. 3(C, VdW balls)). Of particular interest are R346 (BSA: 18.18Å^2^) and T470 (BSA: 21.02Å^2^). This illustrated again the systematic character of MISAs.

## 4 Outlook

The study of biomolecular interactions benefits from experimental and theoretical methods. On the experimental side, structure determination methods provide atomic coordinates for (ensembles) of structures, while thermodynamic/kinetic studies ambition to measure binding affinities and the underlying kinetic properties. Such experiments are often conducted in conjunction with directed mutagenesis, to unveil the specific role of amino acids. On the theoretical side, structure, thermodynamics and kinetics are studied using two classes of methods, namely interaction based methods, and learning based methods. As the name suggest, interaction based methods rely on covalent and non covalent potential energies models and associated force fields. Example such interactions are salt bridges, hydrogen bonds, *π* stacking, etc. Such models generally have a limited spatial and time resolution, which imposes simulation methods to compute observables by means of averages over ensembles. Unfortunately, properties on long time scales are generally out of reach, so that qualitative descriptors of interaction patterns are resorted to. On the other hand, learning based method typically mix geometry based models with biochemical annotations, so as to design proxys for the key determinants of affinity and specificity. Such descriptors are known to be robust, and under suitable hypothesis, have proven good enough to estimate reliably binding affinities.

As illustrated by the example of the RBD of coronaviruses, most studies performed recently tend to promote specific interactions, usually backed up experimentally. This strategy is not entirely satisfactory, and one may actually quote [35] on the interaction (SARS-Cov-1-RBD, ACE2): “ *The residues singled (authors’ note: the 14 a.a.) out for description in the preceding paragraphs are not, of course, the only ones critical for the tight complementarity of the SARS-CoV RBD and human (or palm civet) ACE2.*”. To promote a more comprehensive approach, we introduce MISAs, namely alignments between interface strings defined from geometric models, and coding biological / biophysical properties of interest. While Voronoi interface models are admittedly less precise than interaction based models, they are exhaustive/systematic, parameter free, and can be pulled back onto sequences. The encoding of interactions obtained, called *interface strings*, provides a concise way to capture global and local properties of complexes involving related proteins at a glance, as illustrated while analysis the RBD of coronaviruses.

We anticipate that the encoding of interfaces obtained will become a standard way to concisely represent interactions for families of homologous molecules, helping to understand protein interactions, identify critical residues, and therefore design therapeutics.

## 5 Artwork

## Acknowledgment

T. Dreyfus is acknowledged for insightful comments.

## 6 Supplemental: results

**6.1 Comparing the RBD of SARS-Cov-1 and SARS-Cov-2**

**6.2 Interactions between the RBD of SARS-Cov-2 and antibodies**

**Figure 5:**
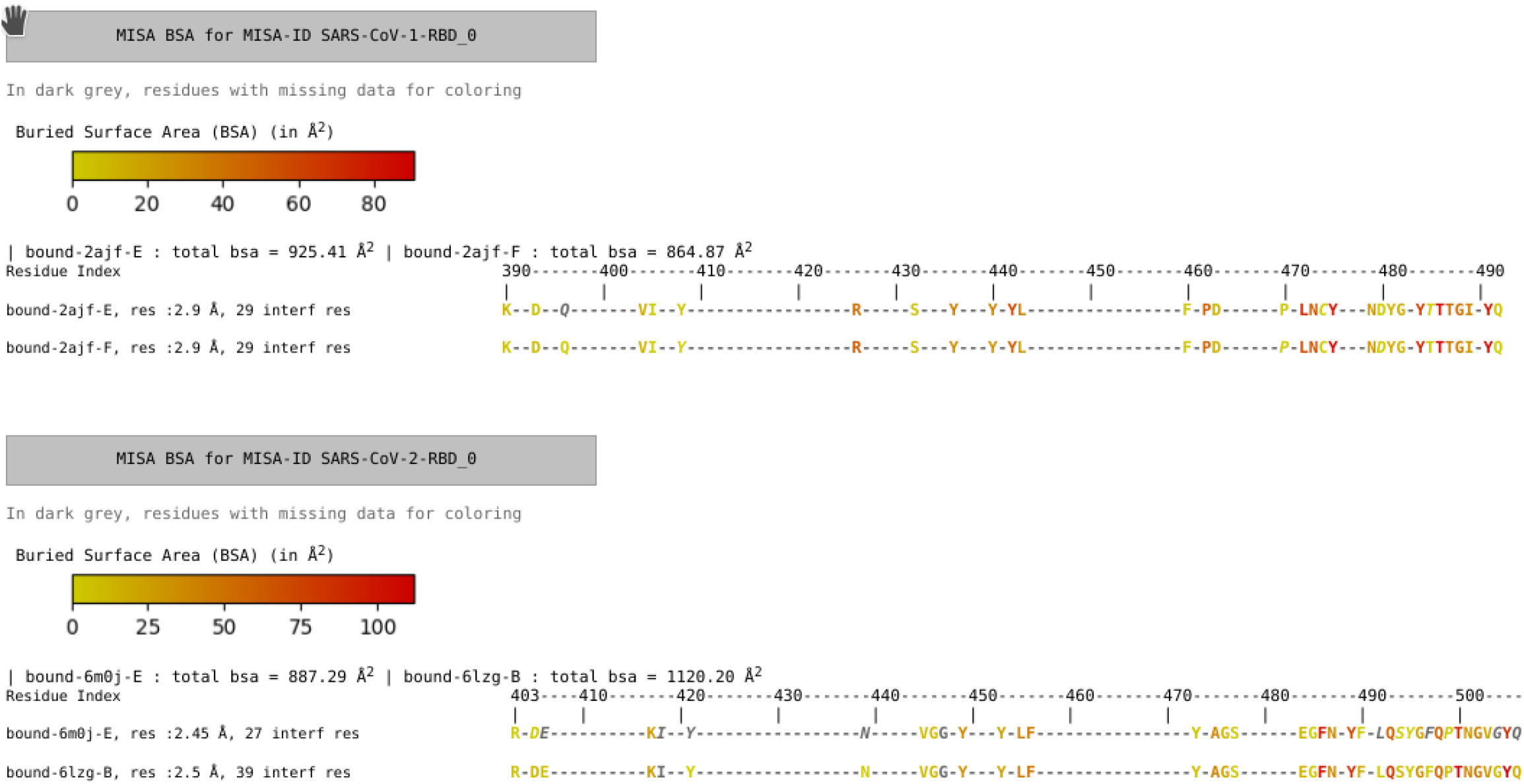
MISA/BSA for Fig. 1.

**Figure 6:**
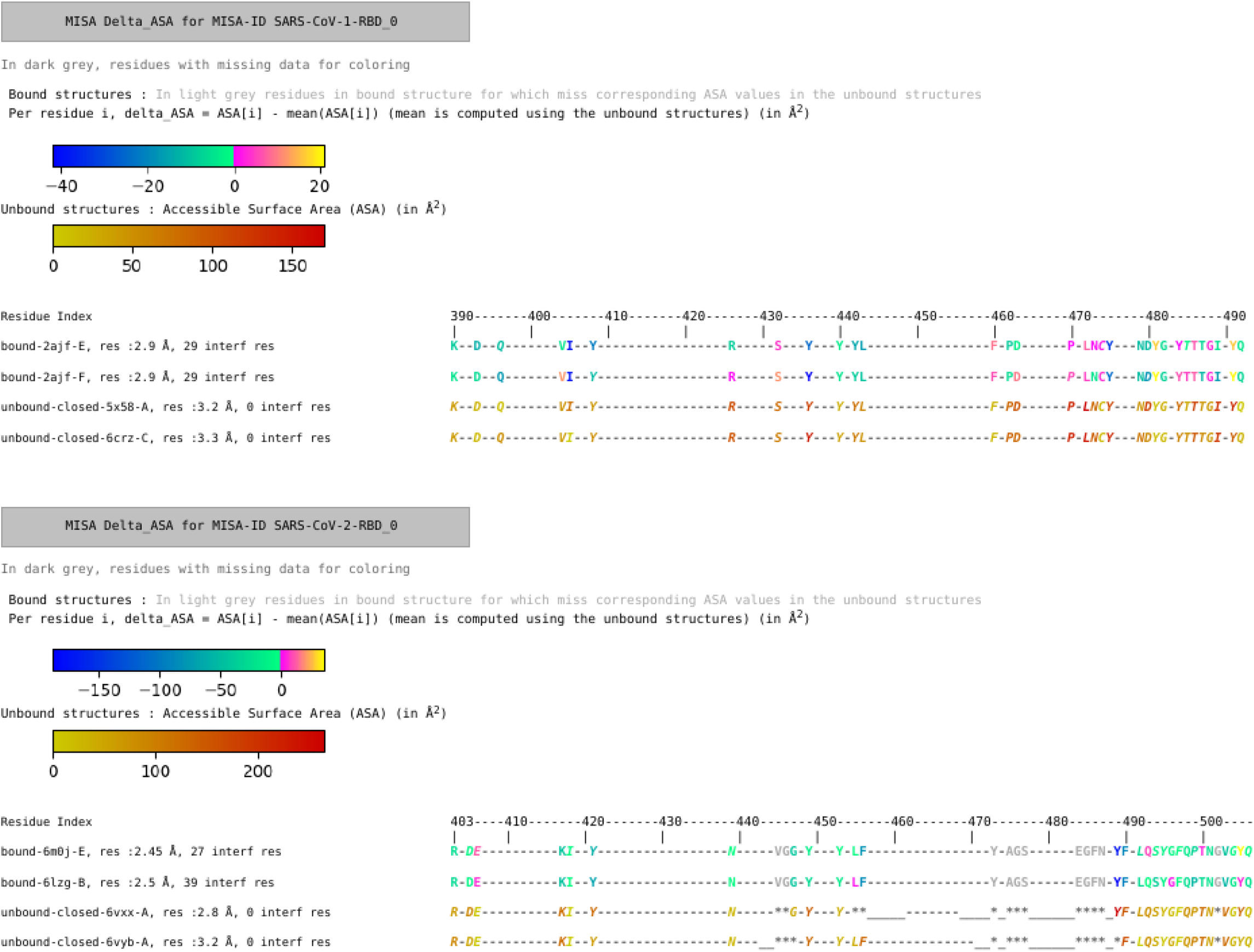
MISA/Δ ASAfor Fig. 1.

**Figure 7:**
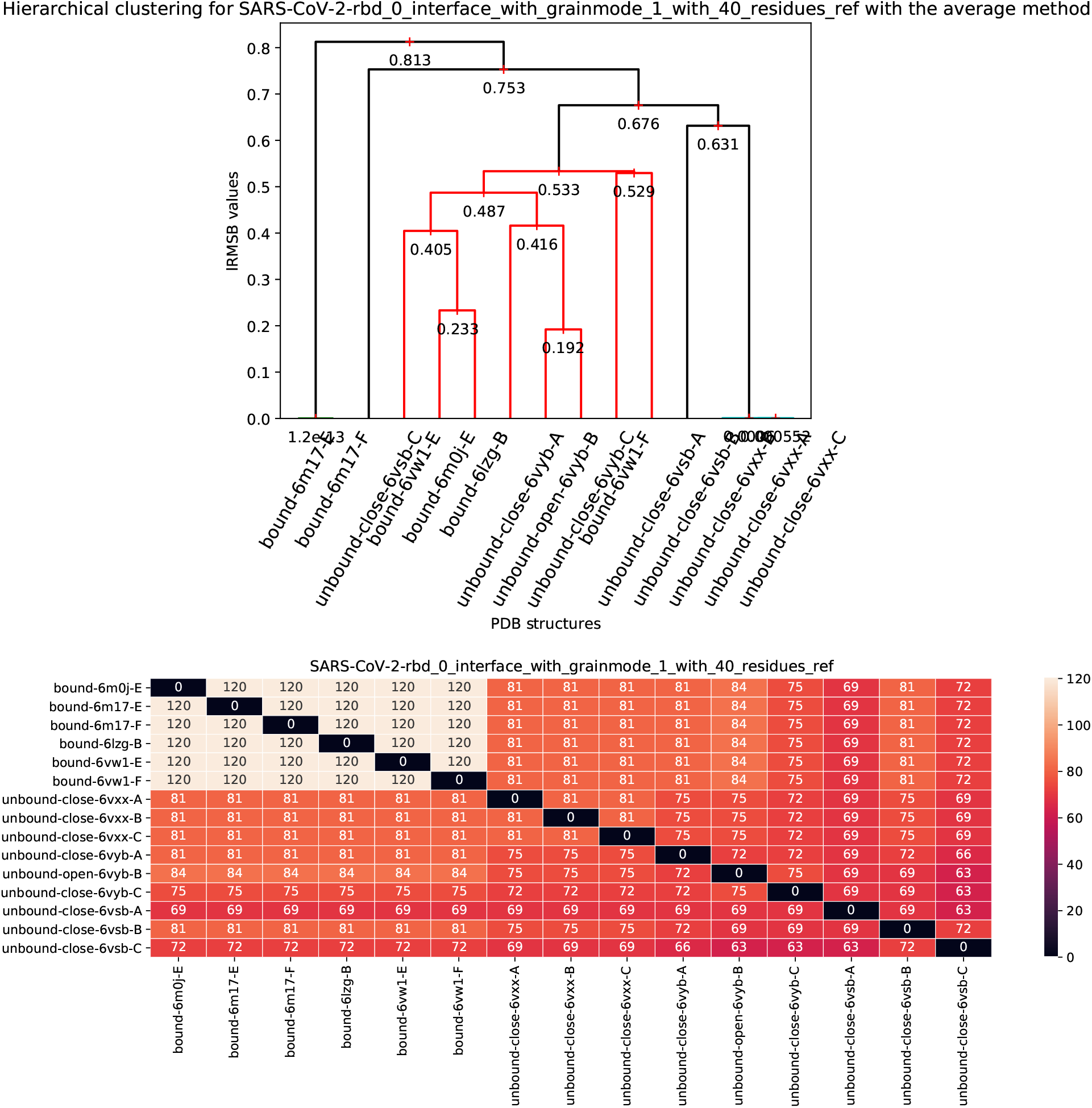
Hierarchical clustering of conformation of the RBD, based on the iRMSD of backbone atoms. Each individual iRMSD is computed based on the backbone atoms common to two structures (selected loops may be missing in a given structure). **(A)** Dendrogram **(B)** Num. backbone atoms in an individual comparison.

**Figure 8:**
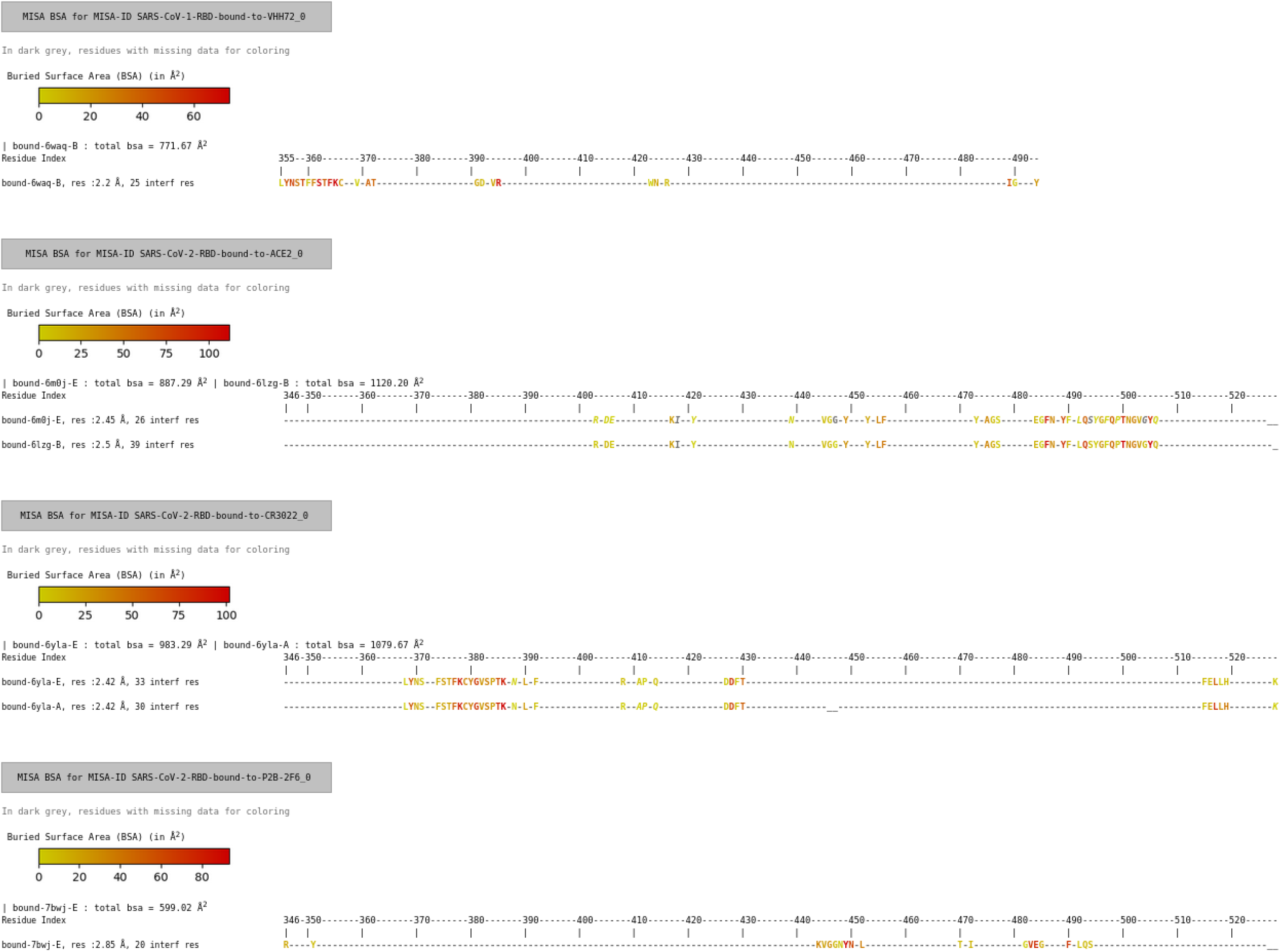
MISA/BSA for Fig. 3.

**Figure 9:**
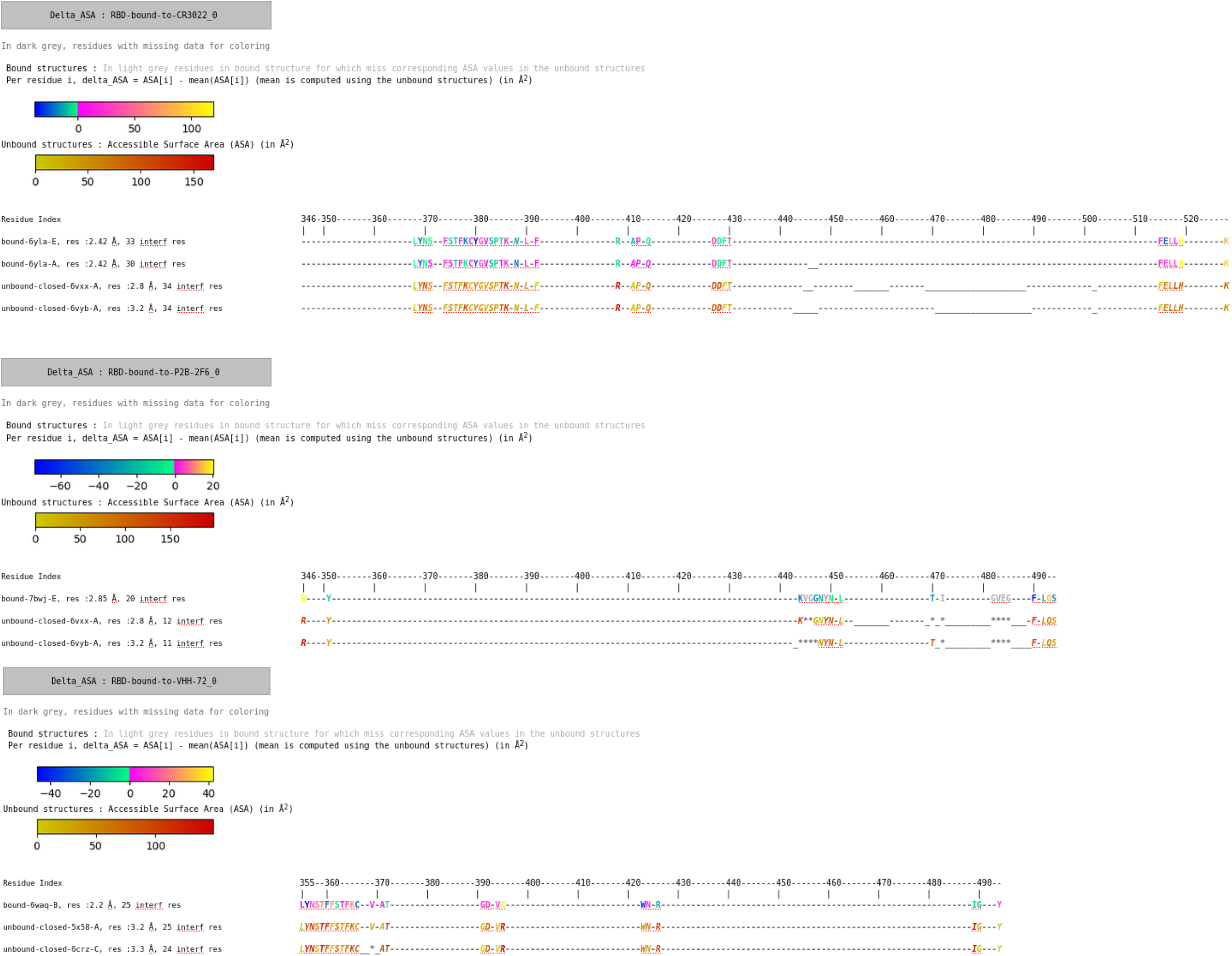
MISA/Δ ASAfor Fig. 3.

## 7 Supplemental: formal specification of the method

We formalize the notions introduced in the main text.

### Voronoi interface restricted to a chain

We first process the Voronoi interface of a complex *C_i_* and collect all the interface amino-acids for a chain instance *S_ij_* = *C_i_*[*A_j_*]:

#### Definition. 5

*(Half interface) Consider the Voronoi interface model for the partners {A_j_} and {B_j_} in structure C_i_. The restriction of the Voronoi interface to chain S_ij_, or* half interface *for short, is the set of a.a. of this chain contributing at least one atom at the interface. This set is denoted 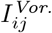. The operator* 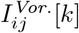 *returns the one letter code of the k-th a.a. of S_ij_ if this a.a. is at the interface, and* ∅ *otherwise.*

Considering all chains associated with a given MISA id yields a set of a.a. at a given position, amidst which we seek the most frequent one:

#### Definition. 6

*The* interface pool *of chain A_j_ at position k is defined as* 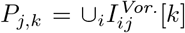, *where index i ranges over all complexes. The* consensus a.a. *at position k is the a.a. most frequent in the pool P_j,k_, with ties broken using the lexicographic on the one letter code of the amino-acids.*

### Interface strings

To encode properties of a particular chain instance *S_ij_* in a structure (bound or unbound), several parameters must be taken into account: the fact that this position is involved in interfaces or not, the potential absence of this position in selected crystal structures, and also the variability observed at that position (Def. 6). These pieces of information are precisely summarized in the so-called interface string, which is a string defined over the alphabet using the one letter code of a.a. plus the three symbols {−_∗_}:

#### Definition. 7

*(Interface string) The* interface string 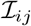 *of chain A_j_ in structure C_i_ or U_i_) is the string with one character per amino-acid, defined as follows:*

- *(Case 1) k-th a.a. never at the interface* i.e. *P_j,k_* = ∅ *: _ if this a.a. is absent from the crystal structure, and - otherwise.*
- *(Case 2) k-th a.a. sometimes at the interface, but not in S_ij_* i.e. *P_j,k_* = ∅ *and* 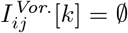: *italicized capital one letter code X for the consensus a.a., italicized lowercase one letter code x otherwise*.
- *(Case 3) k-th a.a. at the interface* i.e. 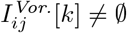:

– *\* if a.a. is absent from the crystal;*
– *capital one letter code* X *for the consensus a.a., lowercase one letter code* x *otherwise.*

Note that for a chain in an unbound structure, Case 3 is irrelevant since there is no interface.

Collecting all such strings for a given MISA id yields Def. 4.

## 8 Implementation details

In the following, we provide the implementation details, using in particular the notations of Section 7 for the operators.

### 8.1 Overview

`sbl-misa.py` is the main script of the package: it creates the MISAs, colors them and performs a structural comparison between the different chains of the same MISA. It uses different modules that we present here, through their main classes.

From its output, different complementary scripts can be used to further the analysis and/or parameterize the display of the MISAs. This section follows the organization of `sbl-misa.py`, and the different steps of the script refer to their numbering in Fig 10.

**Figure 10:**
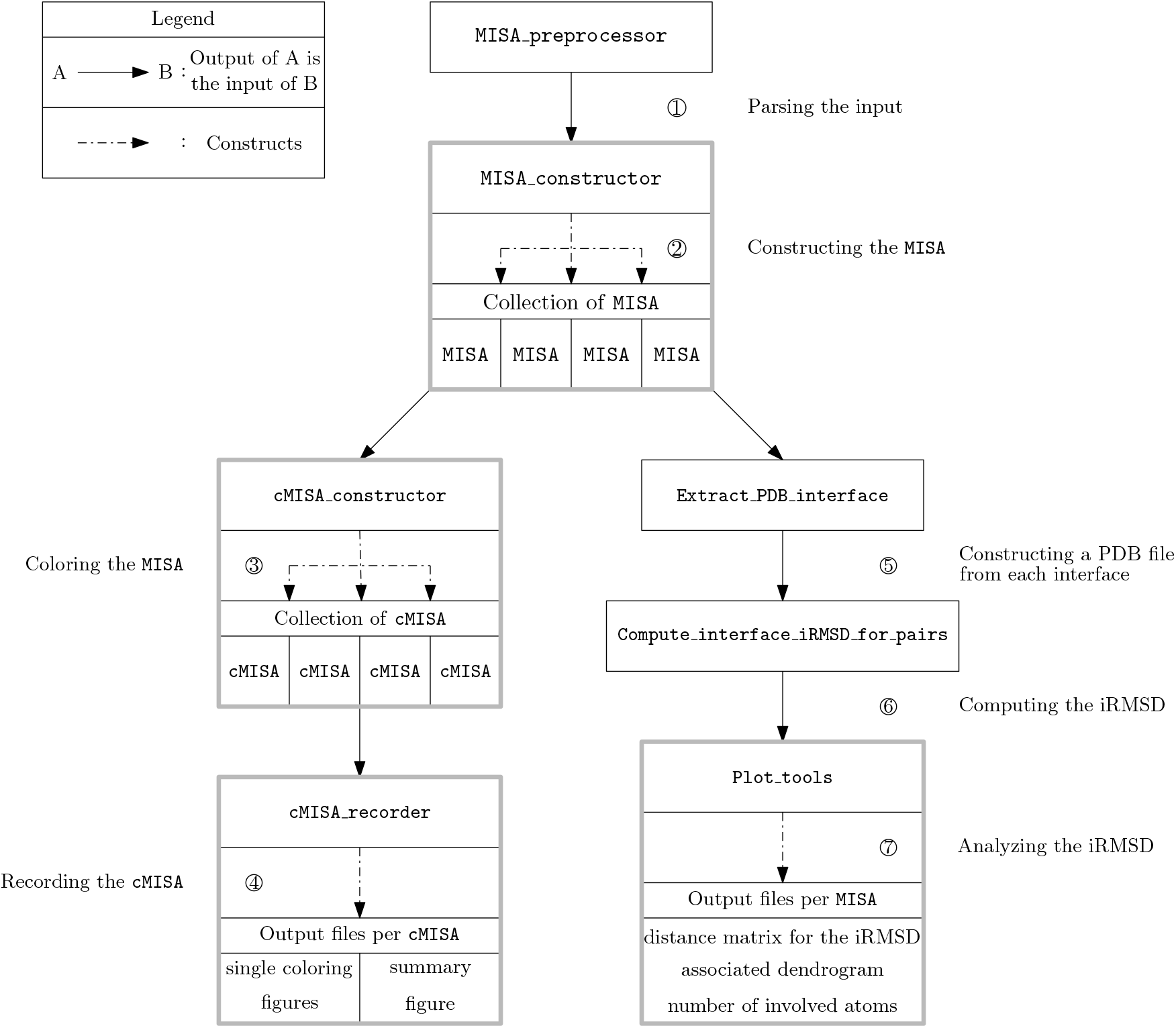
Overview of sbl-misa.py.

### 8.2 Computing the MISA: script sbl-misa.py, module Multiple_interface_string_alignment

#### Overview

`Multiple_interface_string_alignment.py` creates the MISA from a specification file and the input data described above. First, it gathers the homologous structures together, by giving them the same MISA id, and further computes from this set of chains several operators 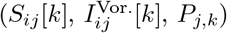. These operators, as presented above, depend on two indices: *i* which designates the complex index and *j* which designates the MISA id. In the code, these operators are implemented at constant *j*, i.e. there is one instance of each operator by MISA id. These operators are used to compute the consensus interface, and from there to create an *istring* for each chain. The simultaneous display of all the *istring* corresponding to the same MISA id is a MISA.

#### Main steps and associated classes

##### ▸ Step 1.: Parsing the input

Main class: **class** `MISA_preprocessor`

*Parse the specification file and gather the structural data*

`sbl-misa.py` starts by reading the input files, and possibly runs `sbl-intervor-ABW-atomic.exe` and/or `sbl-vorlume-pdb.exe`. (The script controls that every structure has a coherent number of chains among the different complex specifications, and if it is not the case, it runs only on the complete structures). It parses the input specification file, and infers the MISA id of each chain.

##### ▸ Step 2.: Constructing the MISA

Main class: **classes** `MISA_constructor, MISA`

*Initialize the MISA by gathering the chains with the same MISA id and aligning them*

In the following, we assume a mapping between the original indexing of the residues in each chain, and the renumbering in the MSA. For homologous chains, this mapping may be computed `ClustalOmega`. The construction of the MISA involves three steps:

- **Initializating the MISA, class** `MISA_constructor`: All the chains sharing the same MISA id are gathered into an instance of the class MISA, which itself contains one instance of the Chain_instance operator *S_ij_*[*k*], one instance of the Interface_oneside operator 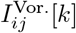 and one instance of the Interface_string operator 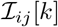 (cf. Fig 2). The classes implementing these three operators have the corresponding names (`Chain_instances, Interface_oneside, Interface_strings`).
- **Constructing the consensus interface, class** `Interface_pool`: The consensus interface is computed for each MISA id from the set of the bound chains, by taking the most frequent residue at each position.
- **Constructing the i-string, class** `MISA`: for each MISA, we compute its instance of 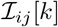, by comparing the residues with the consensus interface and marking them accordingly. Only the residues which are part of a windows, if a windows was provided, are displayed.

###### Remark 3

*Practically, each residue is represented as a triple (aa-or-hetero-code, resid, insertion code), which makes it possible to handle insertion codes. This representation is used both for the keys and values of the aforementioned mapping. The corresponding class is `Residue_mapping`.*

### 8.3 Coloring the MISA: script sbl-misa.py, modules Colored_MISA.py, MISA_ColorEngines.py

#### Overview

`sbl-misa.py` produces four colorings for each `MISA`, one for each of the following quantities (designated as *coloring values*):

- the secondary structure (*SSE*)
- the buried surface area (*BSA*)
- the difference between the accessible surface area (*ASA*) when changing from unbound to bound structure (Δ_*ASA*)
- the B-factor

**Figure 11:**
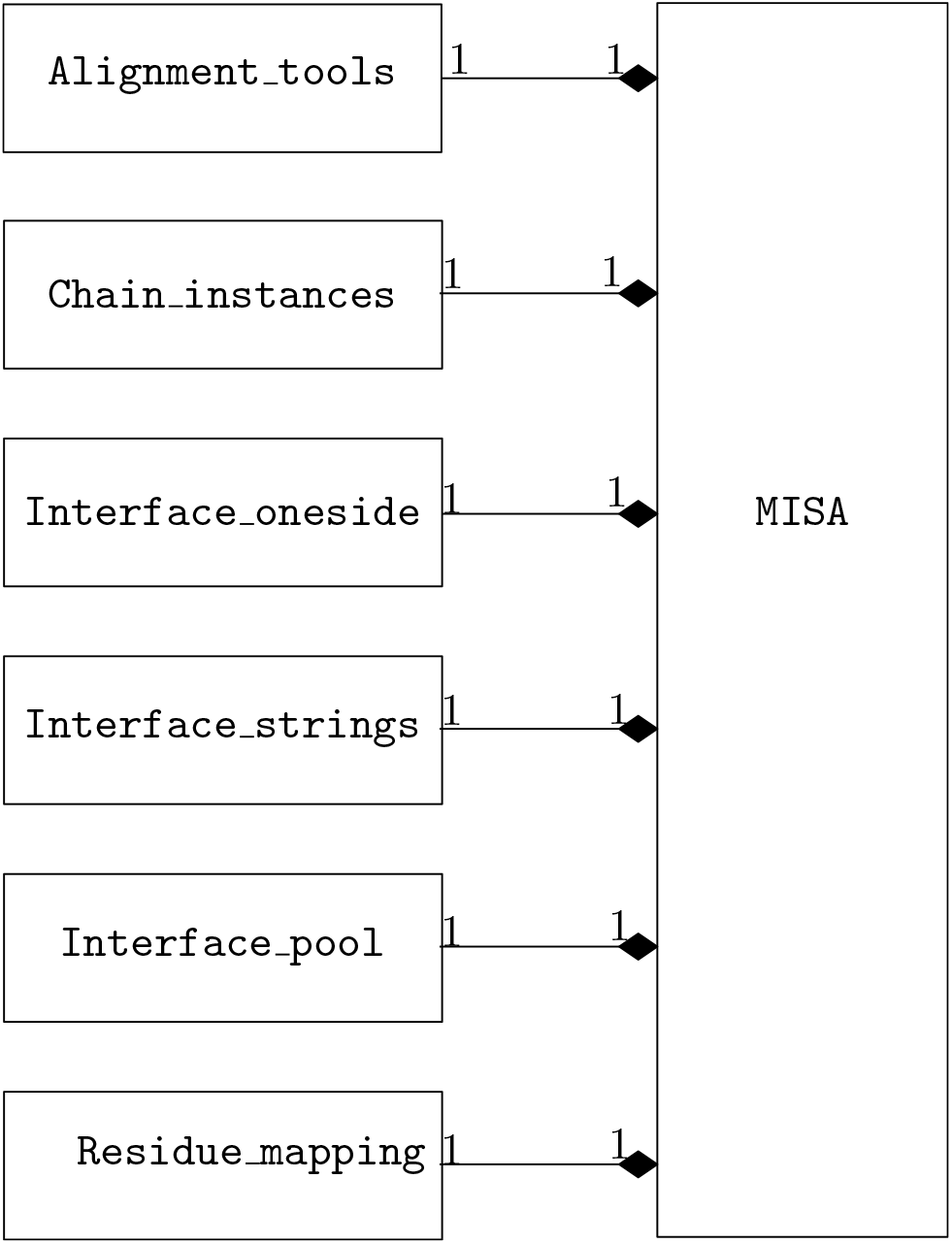
Composition of the MISA class.

**Figure 12:**
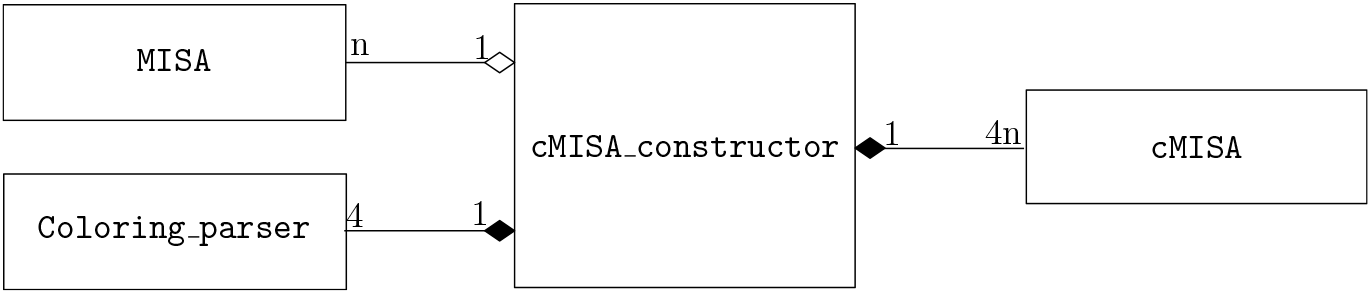
Composition of the cMISA_constructor class.

**Figure 13:**
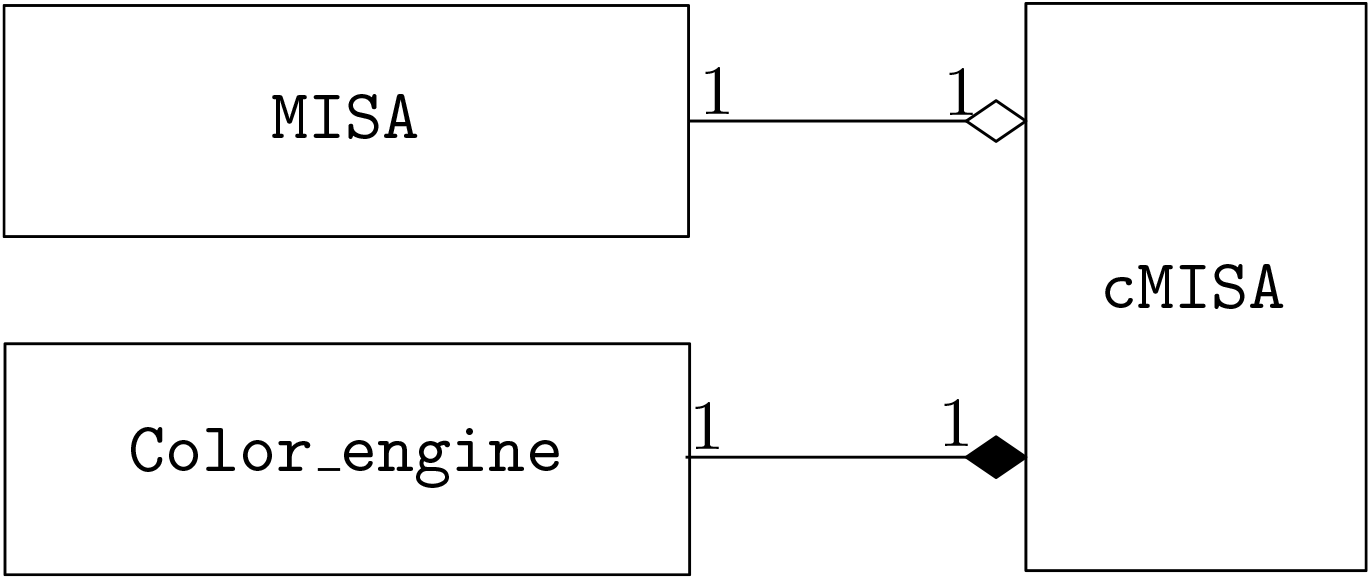
Composition of the cMISA class.

#### Main steps and associated classes

##### ▸ Step 3.: Coloring the MISA

Main class: **classes** `cMISA_constructor, coloring_parser, color_engine`

*Collect the coloring values and attribute the coloring to each chain of the MISA*

- **Collect of the coloring values, classes** `SSE_coloring_parser, BSA_coloring_parser, Delta_ASA_coloring_pars B_factor_coloring_parser`: The coloring values for all the chains are collected by the four coloring_parsers.
- **Creation of the Colored MISA (**`cMISA`**), class** `cMISA_constructor`: For each `MISA`, the `cMISA_constructor` initializes one color_engine per coloring_parser, and creates one instance of the class `cMISA` per pair of `MISA - color_engine`. (Thus we have #`cMISA` = #`MISA` X #`color_engine, with` #`color_engine` = 4) (cf. Fig 3)
- **Application of the coloring, classes `SSE_color_engine, BSA_color_engine, Delta_ASA_color_engine, B_factor_color_engine`**: In each `cMISA` (cf. Fig 3), the color_engine assigns to each of the residues of the `MISA` its *coloring value*, and translates it by the corresponding color, by updating its `Interface_string`.

##### ▸ Step 4.: Recording the cMISA

Main class: **class** `cMISA_recorder`

*Record the colored MISA into an HTML file presenting simultaneously the four colorings*

The `cMISA_recorder` creates the output files, by combining the `Interface_string` of each `cMISA` with the coloring legend provided by the color_engine of the `cMISA`. For each `cMISA`, the chains with the same output are grouped together to lighten the display. A `.html` file displaying simultaneously all the `cMISA` sharing the same MISA id is created in the `odir/MISA` directory. A sub-directory `odir/MISA/single-coloring-figures` is created, where each of the `cMISA` is stored individually, in both `.pdf` and `.html` format.

#### 8.3.1 Coloring SSE

As explained in Section *Colored MISAs*, we color aa in interface strings to show changes in the H bonding network. To this end, we use DSSP [15] and https://swift.cmbi.umcn.nl/gv/dssp/. The corresponding color code reads as follows:

- 3-turn helix: light orange
- 4-turn helix: deep orange
- 5-turn helix: red
- Isolated beta-bridge residue: light blue
- Extended strand: deep blue
- Bend: light green
- Hydrogen bonded turn: deep green
- Other: brown
- Missing Residue: grey

#### 8.3.2 Coloring BSA and Δ_*ASA*

The BSA is computed using the intervor “-bsa” option. For this coloring mode, only bound structures are displayed, as it would make no sense to do so for unbound structures, which necessarily have no BSA.

The ASA is computed using `sbl-vorlume-pdb.exe`, and the Δ_*ASA* is deduced from it, as explained in the main text. We display thus the Δ_*ASA* for the bound structures, and the ASA for the unbound structures. (If only bound structures were provided, the Δ_*ASA* won’t be computable, and the sequence will be displayed in grey).

We use one color map for the BSA coloring, and two for the Δ_*ASA* coloring.

#### 8.3.3 Coloring B-factor

The B-factor is extracted from the PDB file (and possibly computed from ANISOU, see main text, if only ANISOU is available). We use one color map for the B-factor coloring.

### 8.4 Script sbl-misa-mix.py

#### Overview

The `sbl-misa-mix.py` script allows, based on the individual colored MISA produced by `sbl-misa.py`, to display together several colored MISA for an easier comparison.

#### Main steps

- Parse the `mix_ifile`
- Parse the several `.html` files containing the MISA id / colorings of interest
- Combine them into a mixed fig

### 8.5 Script sbl-misa-bsa.py

#### Overview

`sbl-misa-bsa.py` provides access to the numerical values of the buried surface area (BSA) at the residue level, by parsing the .xml file produced by sbl-intervor-ABW-atomic.exe that contains the BSA data.

#### Main steps

- Parse the `spec_file`
- Parse the `xml` file
- Extract the BSA from the appropriate residues

### 8.6 Script sbl-misa-diff.py

#### Overview

`sbl-misa-diff.py` allows to compare the interface of two homologous chains, by comparing the list of their residues at the interface.

#### Main steps

- Parse the `spec_file`
- Gather the data for the different interfaces, either in parsing an `interface-file`, or the MISA output.
- Compare the two interfaces

### 8.7 Structural comparison, script sbl-misa.py, module iRMSD.py

#### Overview

`sbl-misa.py` finally also computes the i-RMSD for each pair of chains within each MISA, to compare the geometry of the different homologous chains.

#### Main steps and associated classes

##### ▸ Step 5.: Construction of the PDB interface files

Main class: **class** `Extract_PDB_interface`

*Store the interface residues shared by each pair of chain into PDB files*

Creates for each pair of chain of each MISA two new `.pdb` files (one per chain), which only contains the interface residues shared by both chains. The selection of the residues at the interface is done thanks to the helper class `ResSelector`.

##### ▸ Step 6.: Computation of the i-RMSD for each pair of chain

Main class: **class** `Compute_interface_iRMSD_for_pairs`

*Compute the iRMSD for each pair of PDB interface files*

Compute the iRMSD for each pair of chains by calling the script `sbl-lrmsd-for-pdb-pair.exe` for each pair of .pdb files precedently created. The l-RMSD is computed for three different ‘grainmodes’ (0: only the C-alpha, 1: the backbone, 2: all the atoms)

##### ▸ Step 7.: Analysis of the i-RMSD

Main class: **class** `Plot_tools`

*Cluster the chains according to the iRMSD and record statistics on the iRMSD*

Once we have the i-RMSD for each MISA, we derive three figures from it:

- The first is a dendrogram showing the hierarchical clustering of the i-RMSD.
- The second is a distance matrix showing the value of the i-RMSD for each pair of chains.
- The third is a matrix showing the number of atoms involved in the calculation of i-RMSD for each pair of chains.

As the i-RMSD is computed for each of the three grainmodes above, these three figures are produced for each of them, and the grainmode is indicated in their title.

